# CellPhenoX: An eXplainable Cell-specific machine learning method to predict clinical Phenotypes using single-cell multi-omics

**DOI:** 10.1101/2025.01.24.634132

**Authors:** Jade Young, Jun Inamo, Zachary Caterer, Revanth Krishna, Fan Zhang

## Abstract

Single-cell technologies have enhanced our knowledge of molecular and cellular heterogeneity underlying disease. As the scale of single-cell datasets expands, linking cell-level phenotypic alterations with clinical outcomes becomes increasingly challenging. To address this, we introduce CellPhenoX, an eXplainable machine learning method to identify cell-specific phenotypes that influence clinical outcomes. CellPhenoX integrates classification models, explainable AI techniques, and a statistical framework to generate interpretable, cell-specific scores that uncover cell populations associated with relevant clinical phenotypes and interaction effects. We demonstrated the performance of CellPhenoX across diverse single-cell designs, including simulations, binary disease-control comparisons, and multi-class studies. Notably, CellPhenoX identified an activated monocyte phenotype in COVID-19, with expansion correlated with disease severity after adjusting for covariates and interactive effects. It also uncovered an inflammation-associated gradient in fibroblasts from ulcerative colitis. We anticipate that CellPhenoX holds the potential to detect clinically relevant phenotypic changes in single-cell data with multiple sources of variation, paving the way for translating single-cell findings into clinical impact.

## INTRODUCTION

Single-cell omics technologies have revolutionized our understanding of biological heterogeneity, and offer an opportunity to identify novel cell phenotypes potentially linked to disease pathogenesis1. With the exponential growth of single-cell data generating billions of cell profiles through collaborative consortia efforts to unravel biological complexity2, a new computational challenge has emerged: how to reliably detect phenotypic changes associated with clinically relevant attributes3. Building on early approaches that used clustering for differential abundance testing4,5, recent methods have evolved to address this association problem in a cluster-free manner that provides finer granularity in identifying outcome-associated cells compared to cluster-based methods6. Most of the cluster-free methods, such as Meld7, Milo8, and CNA9, construct transcriptional neighborhoods to identify cell neighbors associated with sample-level outcomes. A benchmarking study further demonstrated that Milo effectively maintained accuracy, particularly in the presence of substantial batch effects, while CNA provided scalability for large single-cell datasets6. However, these methods primarily rely on linear mixed-effects modeling to determine statistical associations and lack a machine learning framework to enhance predictive capability. Moreover, they are not equipped to detect cell populations influenced by interaction effects, a phenomenon commonly observed in many diseases.

Given the complexity of large single-cell data in a multi-sample, multi-disease stage, and complex clinical setting, artificial intelligence (AI) techniques have the potential to excel in uncovering hidden patterns to identify robust biomarkers more effectively than traditional statistical or linear models. As AI becomes more prevalent, explainable AI (XAI) strives to enable the users to gain a better understanding of the factors influencing the model’s output10. Techniques such as the SHapley Additive exPlanations (SHAP) values provide clarity on how features influence model predictions, offering insights into both linear and nonlinear interactions11,12. Several applications of XAI are used to identify protein markers for sample-level measurements like plasma proteomics or bulk experiments13. However, it remains unclear whether such XAI techniques are capable of identifying cell-level phenotypic changes that have classifiable and predictable attributes, and whether they are effective for complex disease contexts that often contain confounding factors, such as infectious and immune-mediated diseases14–17.

Here, we introduce CellPhenoX, an eXplainable machine learning tool to identify cell-specific phenotypes that influence clinical outcomes of interest for single-cell studies. CellPhenoX classifies clinical phenotypes while accounting for covariates and interactive effects, generating cell-specific Interpretable Scores to summarize the discriminative power of model prediction. By benchmarking CellPhenoX on simulated and real single-cell disease datasets against other existing methods, we demonstrate its robust calibration in detecting condition-associated cell populations, especially rare cell phenotypes and interaction effects, in both binary disease-control and multi-class clinical outcomes. The interpretable scores revealed by CellPhenoX provide new insights into the existing statistical linear model-based associations, opening new avenues to link molecular and cellular subphenotypes with future clinical impact.

## RESULTS

### Overview of CellPhenoX

The goal of CellPhenoX is to integrate the robustness of classification models with the transparency of XAI methods to produce interpretable, cell-specific scores that identify cell populations that can be used to classify a clinical phenotype of interest (**Figure 1**). First, we transform single-cell gene expression into cell abundance across samples using a neighborhood abundance matrix (NAM), then apply the preferred dimensionality reduction method to the NAM to obtain latent dimensions *Xi*, where *i* represents the number of latent dimensions. To predict the clinical phenotype of interest *Y* (e.g., disease status, treatment response), we use *Xi* as features in our classification models, along with proper covariates *γ* (e.g., technical batch) and doptional interaction effect *δ* (e.g., age). Incorporating these factors explicitly into the model allows us to account for their influence rather than regressing out their effects, thus preserving data integrity and enhancing interpretability. The classification model is trained using a nested cross-validation (CV) strategy to ensure robustness and prevent overfitting. Final model performance is determined using a hold-out validation set, providing an unbiased assessment of predictive capability (**Methods**). Within this framework, we employ Shapley Additive Explanation (SHAP)10 values to quantify the contribution of each feature to the prediction for each cell. SHAP provides a local importance score for each feature, allowing us to assess contributions per individual cell. We then rank features based on the distribution of their mean absolute SHAP values, and generate an Interpretable Score by summing the SHAP values across features for each cell. The Interpretable Score allows us to link annotated cell populations to their associations with the clinical phenotype of interest and project the scores onto a UMAP18 low-dimensional space to facilitate the identification of patterns related to the phenotype. Furthermore, we identify markers whose expression levels are significantly correlated with the Interpretable Score, providing additional biological insights. Since CellPhenoX is designed to model changes in cell abundance based on their predictive or discriminatory capability for clinical metrics, this approach can be generalized to multiple single-cell sequencing modalities by providing interpretable results that link single-cell heterogeneity to clinical impact.

**Figure 1.**
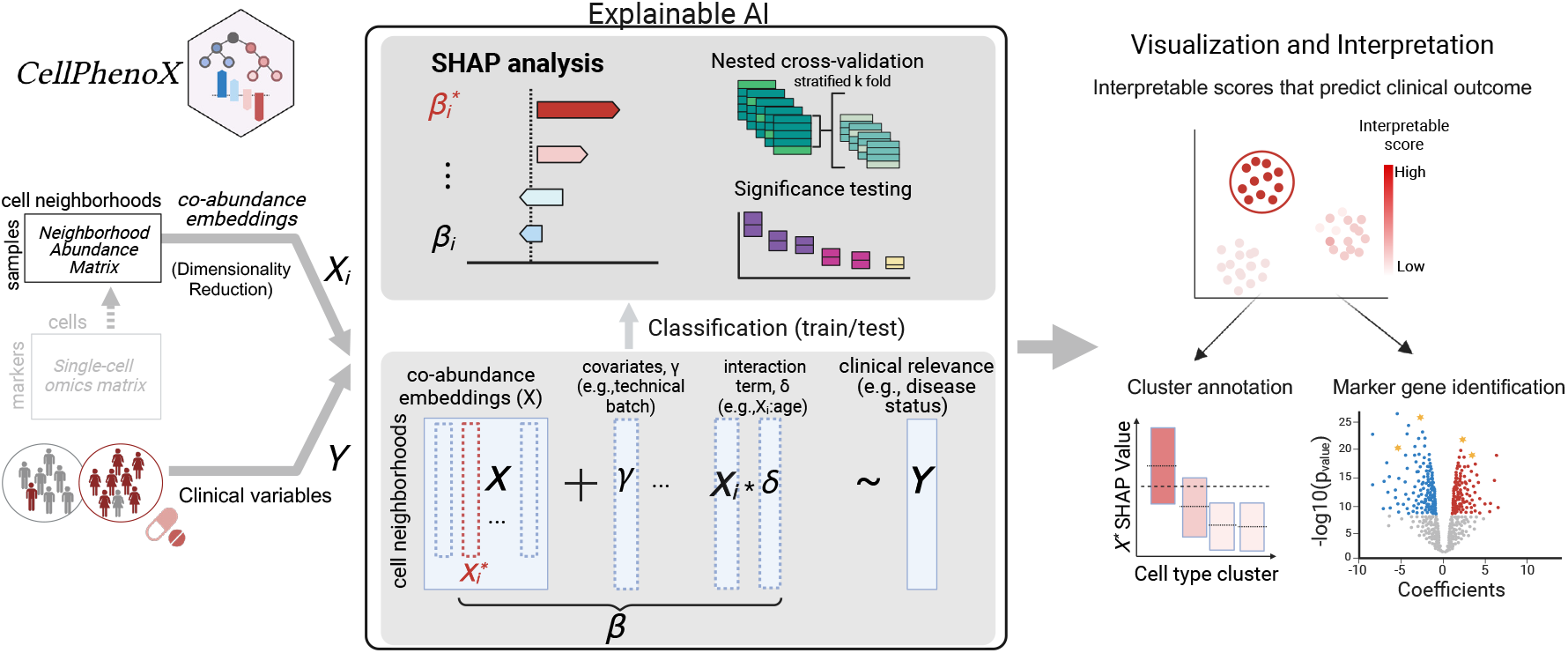
Overview of CellPhenoX methodology. CellPhenoX leverages cell neighborhood co-abundance embeddings, *Xi*, across samples and clinical variable *Y* as inputs. By applying an adapted SHAP framework for classification models, CellPhenoX generates Interpretable Scores that quantify the contribution of each feature *Xi*, along with covariates *γ* and interaction term *Xi* * *δ*, to the prediction of a clinically relevant phenotype *Y*. The results are visualized at single-cell level, showcasing Interpretable Scores at low-dimensional space, correlated cell type annotations, and associated marker genes.

### CellPhenoX demonstrates superior power on single-cell simulation data

To evaluate the performance of CellPhenoX, we designed our simulated data with varying levels of cell cluster abundance in a disease-control setting (**Methods**). We simulated two datasets, each with 15 disease samples and 15 control samples, containing around 100 cells per cell type per sample. Simulation dataset 1 has 10 cell clusters, with cell type clusters A and J expanded in disease compared to control. Specifically, we made cluster J to represent a rare cell type which is more challenging to determine in terms of its disease association. In contrast, Simulation dataset 2 represents a more complex case, where all ten clusters show a slightly higher proportion of disease-associated cells compared to controls. Compared to controls, the fold change ratio of clusters A and J in diseased samples is 3 in dataset 1 and 0.1 in dataset 2, respectively (**Figure 2A, Supplementary Figure 2A**).

**Figure 2.**
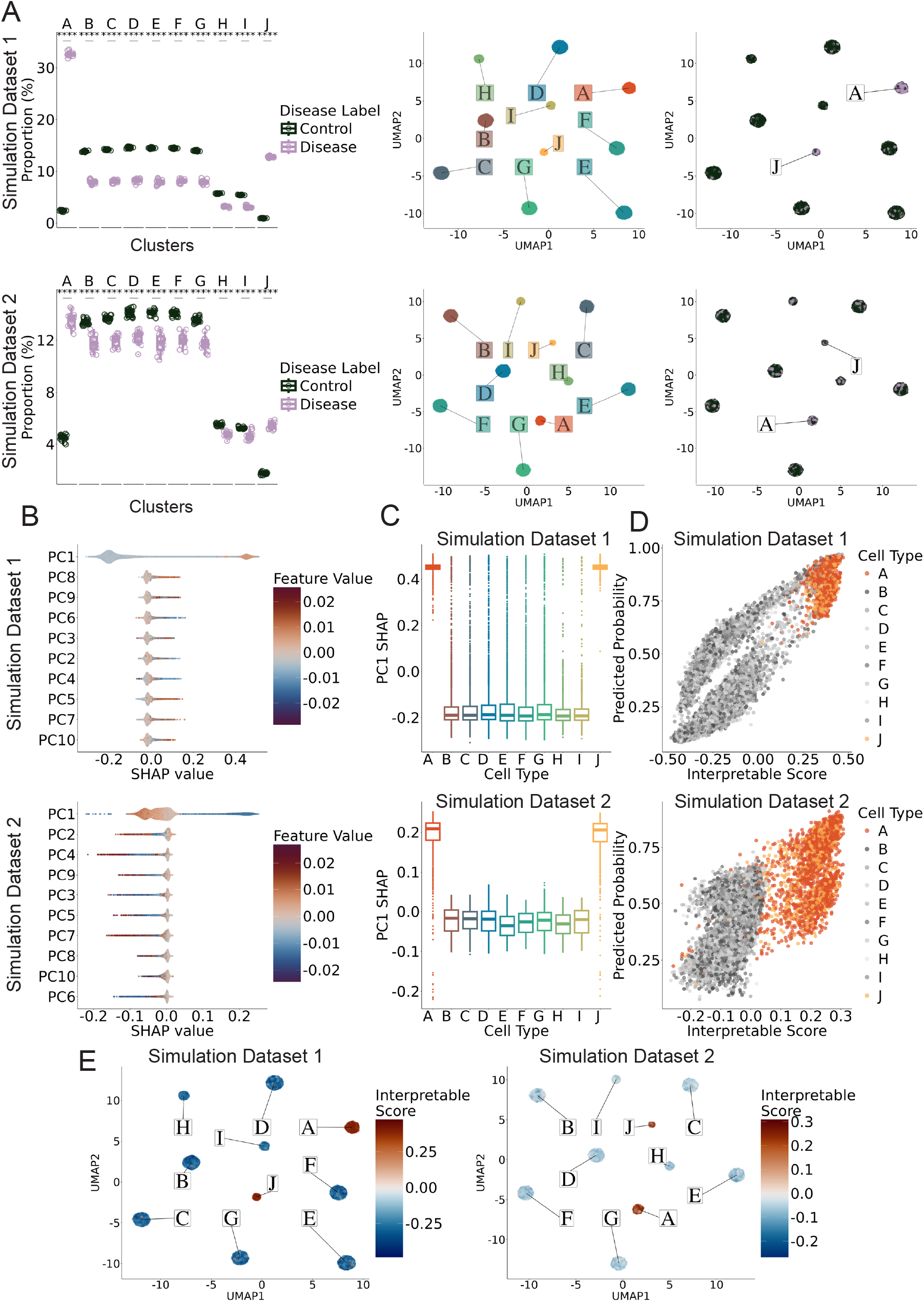
Performance of CellPhenoX on simulation single-cell datasets. **A**. Boxplots depicting the proportion of cells in disease and control groups for each simulated cell type cluster (left), UMAP plot colored by the cell type cluster (middle) and by disease status (right). The top row represents Simulation Dataset 1, where the fold change between the proportions of disease cells and control cells is 3.0. The bottom row corresponds to Simulation Dataset 2, where the fold change between the proportions of disease cells and control cells is 0.1, **B**. SHAP summary plots showing the features ranked by the mean absolute SHAP value where each point represents an individual cell, the color denotes the feature value, and the position along the x-axis represents the corresponding SHAP value, **C**. Boxplots showing the distribution of SHAP values across cell types for the most influential feature, PC1 SHAP, **D**. Scatterplot comparing the discriminatory power of the random forest predicted probabilities and CellphenoX Interpretable Score; here clusters A and J are simulated disease associated cell types, **E**. Visualization of the Interpretable Score indicating disease association on the UMAP plot.

Next, we applied CellPhenoX to each of the simulation datasets by generating a neighborhood abundance matrix (NAM) and decomposing it using PCA. These latent PCs were input into CellPhenoX to predict disease status using Random Forest (**Methods, Supplementary Figure 1B). Under CellPhenoX, we calculated SHAP summary results highlighting the significant influence of PC1 on the model’s predictions (Figure 2B**). Notably, PC1 indeed accurately identified clusters A and J as the drivers of the differences between disease and control samples (**Figure 2C**). Furthermore, we compared the discriminative power of the random forest model’s predicted probabilities with CellPhenoX’s Interpretable Score. The analysis indicates that our Interpretable Score more effectively separates cell types A and J from the rest of the cell types compared to random forest model’s predicted probabilities (**Figure 2D**), especially in Simulation dataset 2. We further presented our CellPhenoX Interpretable Score in the UMAP, confirming that higher scores correspond to cell profiles in the simulated disease expanded clusters A and J (**Figure 2E**).

Additionally, we benchmarked CellPhenoX by comparing its performance with CNA9 and MiloR8, evaluating the results from each method against the true disease status of each simulated cell (**Methods**). We found that the median proportion of correct predictions for CellPhenoX across all simulated clusters is 0.848 for Simulation Dataset 1 and 0.710 for Simulation Dataset 2, which outperformed the results from MiloR and CNA (**Supplementary Figure 2**). While MiloR was able to identify cell type A as expanded in disease, it failed to detect the rare cell type J that was also simulated to be expanded (**Supplementary Figure 2C-D**). Additionally, we observed that miloR results may be unstable depending on the selection of cells used to construct neighborhoods (**Supplementary Figure 2E**). Altogether, CellPhenoX demonstrated outstanding performance in identifying condition-associated cell populations, with superior capability in detecting rare cell populations with precision. While benchmarked methods here reveal statistical correlations relying primarily on linear models, CellPhenoX offers valuable insights into interpretable contribution of individual cells to the predictive outcome.

### CellPhenoX deciphers interaction effects between sex and disease

Disease status rarely manifests in isolation but instead results from an interaction between the condition with other sources of variation, such as sex or age. To our knowledge, there is a need for methods capable of deciphering the interaction effects in single-cell data, as most existing approaches, such as CNA and MiloR, lack the functionality to detect such interactions. In our study, we simulated a single-cell dataset featuring differentially abundant clusters influenced by the interaction between sex and disease. This simulation represents a scenario in which disease status interacts with the covariate sex, affecting two major cell types, A and B, and that are depleted and expanded in female disease, respectively (**Figure 3A**). We also simulated minor cell types I and J, which are also depleted and expanded in female disease, but pose a challenge as they form rare clusters (5%) and also cluster with non-disease associated cell types in UMAP space (**Figure 3A-B**). Applying CellPhenoX, we are able to predict binary disease status using the harmonized NAM PCs, their interaction terms with sex, and sex as a covariate. Notably, the most salient feature we identified was the interaction term between the first latent dimension and sex (**Supplementary Figure 3A**). Specifically, we observed that the Interpretable Score was approximately zero for cells from male individuals, while for female individuals, it was approximately 0.35 for cell types B and J and -0.35 for cell types A and I (**Figure 3C**). This clearly identified the clusters influenced by the interaction effect. We further demonstrated that the CellPhenoX Interpretable Score recaptured disease- and sex-biased signals in UMAP space, pinpointing both major cell type B and minor cell type J with sex-biased disease preferences (**Figure 3D**). These findings highlight that CellPhenoX is capable of extracting fine-grained disease signals arising from complex interactions between cellular components and covariates.

**Figure 3.**
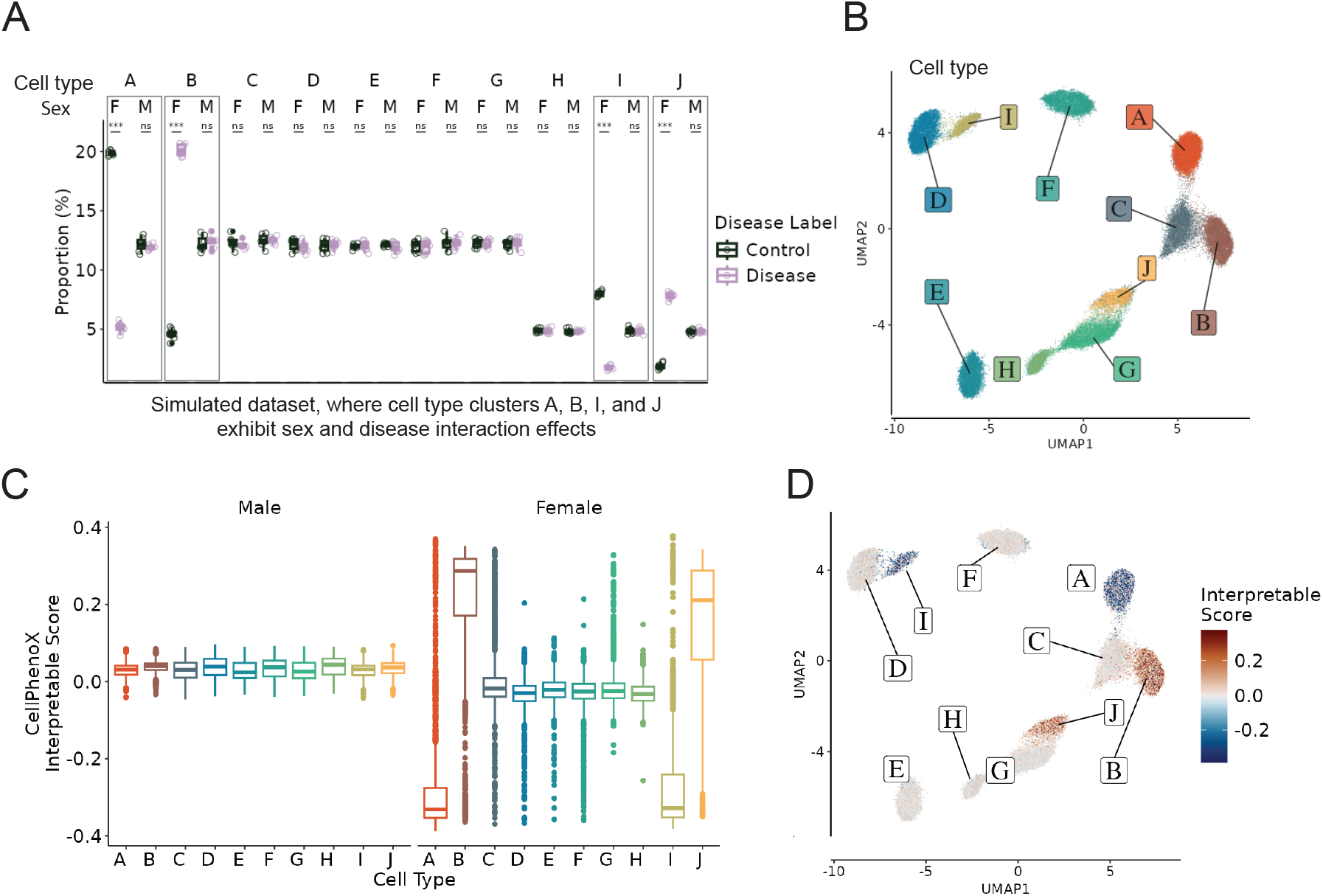
CellPhenoX identifies cell types influenced by disease and sex interactions in simulated single-cell dataset. **A**. Boxplots showing the proportion of cells in disease and control groups in Simulation Dataset for each simulated cell type and sex, where “M” is male and “F” is female, **B**. UMAP plot colored by cell type clusters, **C**. Boxplots faceted by sex showing the distribution of the CellPhenoX Interpretable Scores for each cell type, **D**. Visualization of the Interpretable Score indicating sex-predominant disease associations on the UMAP plot.

### CellPhenoX identifies an activated macrophage phenotype in COVID-19 using single-cell proteomics

To assess the potential of CellPhenoX to identify specific cell phenotypes implicated in disease, we applied it to a large COVID-19 proteomics dataset16. Using CellPhenoX, we performed the multi-class classification of healthy, mild COVID-19, and moderate COVID-19 stages to elucidate cells associated with increasing disease severity. Preliminary investigation of cell density across these conditions revealed an increased abundance of the myeloid cells, NK cells, and CD8+ T cells (**Figure 4A**). However, this observation is only driven by cell density and does not account for technical effects, sample variation, or key demographic variables such as age and sex. To accurately capture disease-associated abundance shifts with classification robustness, we are able to predict disease status-associated abundance changes reflected in the latent components, specifically NAM PCs, adjusting for sex, age, days from disease onset, and smoking status, in our cross-validation interpretable classification model. Note that the NAM PCs are batch corrected using Harmony19 to reduce the technical batch effect. Furthermore, we included age and sex as interaction terms into our model to capture the complex relationships between these covariates and the latent components (**Methods**). To ensure consistency in providing one Interpretable value for each cell, we aggregated the SHAP values from the three disease statuses as described in the multi-class SHAP aggregation Methods section (**Figure 4B**). Thus, according to the original cell type annotations16 (**Figure 4C left**), CellPhenoX’s interpretable score highlighted the myeloid cell population as the strongest predictor for the moderate stage (**Figure 4C right**). The summary plot of the ten most influential latent PCs further underscores the importance of the low-dimensional cell component interaction terms compared to the standalone cell components, with the interaction of NAM PC1 and age contributing the most to the model prediction (**Figure 4D**). We further demonstrated the commutative property of SHAP values by decomposing our Interpretable scores and mapping them to cell type annotations. Notably, proliferating monocytes, CD14+, and CD16+ monocytes exhibited a more distinct signal as subsequent features were included in the model (**Figure 4E**). This conveys the robustness of CellPhenoX’s Interpretable Score which consistently identifies proliferating monocytes—a minor but important myeloid cell phenotype for COVID-19.

**Figure 4.**
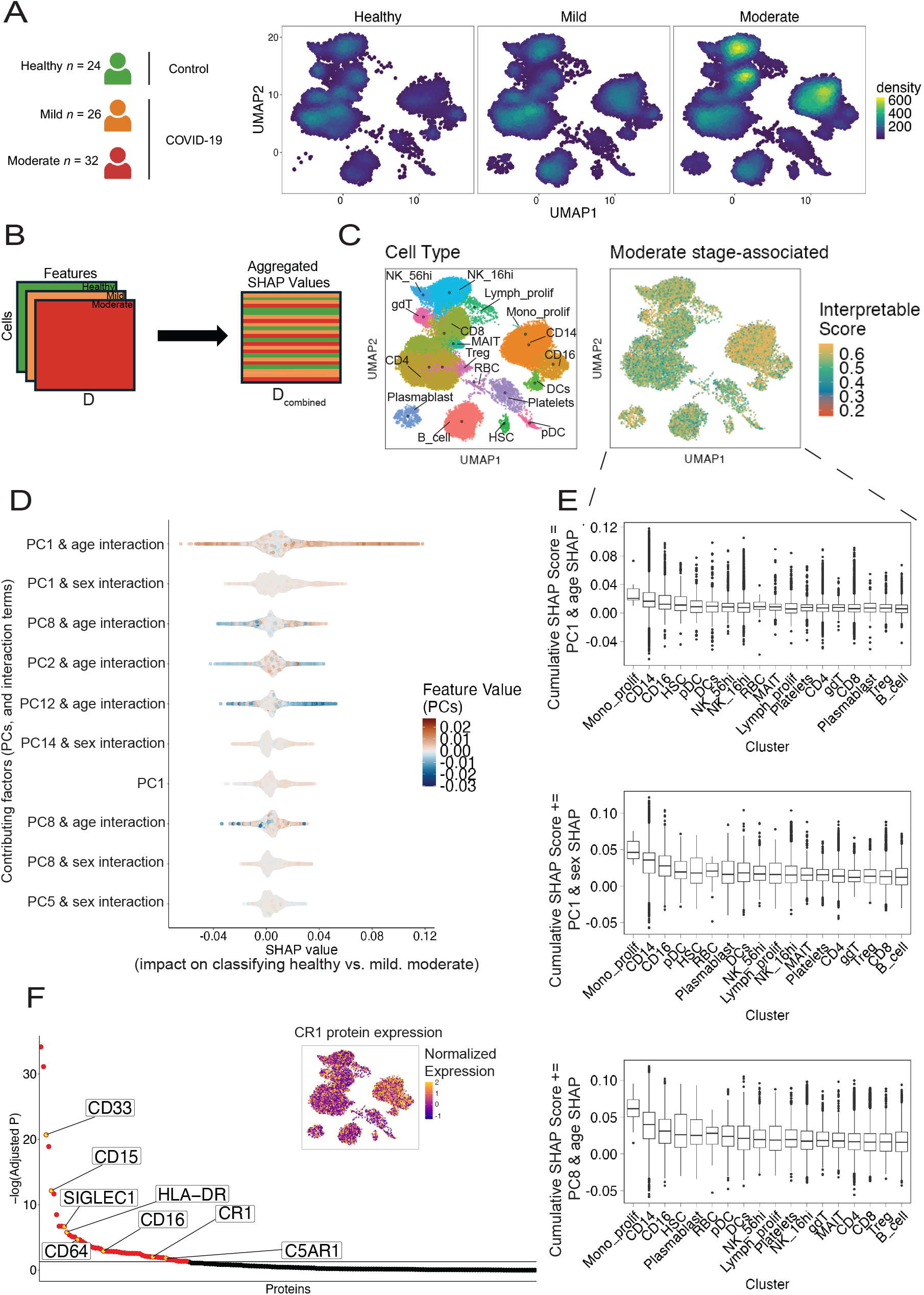
Applying CellPhenoX to single-cell proteomics of PBMCs from a COVID-19 study. **A**. A schematic plot of the multi-class design including the samples from healthy, mild, and moderate COVID-19 severity classes (left), and UMAP plots for cell density for each healthy, mild, and moderate classes, **B**. Depiction of the aggregation procedure where the SHAP values for each cell are selected from the SHAP value matrix that corresponds to the prediction for that cell (**Methods**), **C**. UMAP plot annotated with the cell cluster labels from the original study (left) and UMAP colored by the Interpretable score by CellPhenoX (right), **D**. SHAP summary plot showing the top 10 features ranked by the mean absolute SHAP value, where each point represents an individual cell, the color denotes the feature value, and the position along the x-axis represents the corresponding SHAP value, **E**. Boxplots showing the distribution of CellPhenoX’s Interpretable Score for each cluster starting with the most salient feature (top), then the cumulative SHAP score with the second (middle) and the third (bottom) most influential feature added, **F**. Proteins sorted by adjusted p-value regarding their correlations with Interpretable Score. Yellow points highlight specific proteins that are related specifically to monocyte lineage for COVID-19 severity by literature. CR1 protein normalized expression is presented.

The generated Interpretable Score can be used to identify key associated molecular markers. We next examined the proteins with expression levels significantly correlated with the generated Interpretable Score for cells from moderate patients (adjusted *p*<0.05). In turn, we elucidated key protein markers associated with the discriminative capability of the Interpretable Score, particularly CD33, CD15, SIGLEC1, HLA-DR, CD64, CR1, C5AR1, and CD16 (**Figure 4F**).

Among these, SIGLEC1 was reported to be expressed by circulating monocytes in COVID-19 and associated with disease severity20. CD33 was reported as a novel biomarker in peripheral monocytes for COVID-19 severity21. Further, key immune complement components, including C5AR1 and CR1 (**Figure 4F)**, were uncovered, which were recently shown to increase with COVID-19 severity leading to the activation of myeloid cells22,23. Further investigation is needed to evaluate the potential of these serving as a target for halting disease progression. Overall, these findings highlight the robustness of CellPhenoX in identifying cell phenotypes and relevant markers associated with changes in disease severity, while accounting for critical covariates and interaction effects.

### CellPhenoX reveals subtle fibroblast-specific phenotypic changes unique to in inflamed ulcerative colitis using single-cell transcriptomics

It is crucial to identify cell type-specific variations that are associated with clinical relevance, but this task is often computationally challenging due to the high heterogeneity, especially in diseased tissues. To evaluate CellPhenoX on a specific cell type, we applied it to the single-cell transcriptomics of fibroblasts from patients with Ulcerative Colitis (UC)24. In this study, adjacent inflamed and non-inflamed biopsies were collected from patients with UC (n=18; **Figure 5A**). To uncover the biological topics underlying cell co-abundance, we used NMF25, a topic modeling method, and trained a random forest model to predict inflamed vs non-inflamed status (**Methods**), yielding a classification AUROC of 0.8, AUPRC of 0.86 (**Supplementary Figure 4A**). We observed that NMF-derived latent dimensions 1 and 3 presented the most influence on the model prediction (**Figure 5B**). Based on the fibroblast cluster annotations from original study24, we found that the inflammatory fibroblast cluster exhibited the highest Interpretable Score on average (*p*<2.2e-16), implicating its critical role in the progression of tissue inflammation (**Figure 5C**). This is further confirmed by visualizing the cell-specific Interpretable Score, where inflammatory fibroblasts consistently show the highest scores, with a clear gradient extending toward a subset of *WNT2B+* fibroblasts (**Figure 5E**). Furthermore, we determined the genes that are correlated with the discriminatory power of the Interpretable Score (**Methods**). Among the 21 significant genes (adjusted *p*<0.05) (**Figure 5F**), we identified that *WNT2B+* marker genes (e.g., *ADAMDEC1, CXCL12, CFD*, and *APOD*) are negatively associated with Interpretable Score, while inflammatory fibroblast marker genes (e.g., *CCL19, CCL8, COL6A3*, and *IGFBP5)* are positively associated with the Interpretable Score (**Figure 5G**). This indicates that the continuous state transition from *WNT2B+* subset to inflammatory fibroblast subset may serve as a key predictor of tissue inflammation, warranting further investigation. In addition, for model robustness, we also tested a different dimensionality reduction method, PCA, which yielded a similar inflammatory fibroblast axis (**Supplementary Figure 4B**).

**Figure 5.**
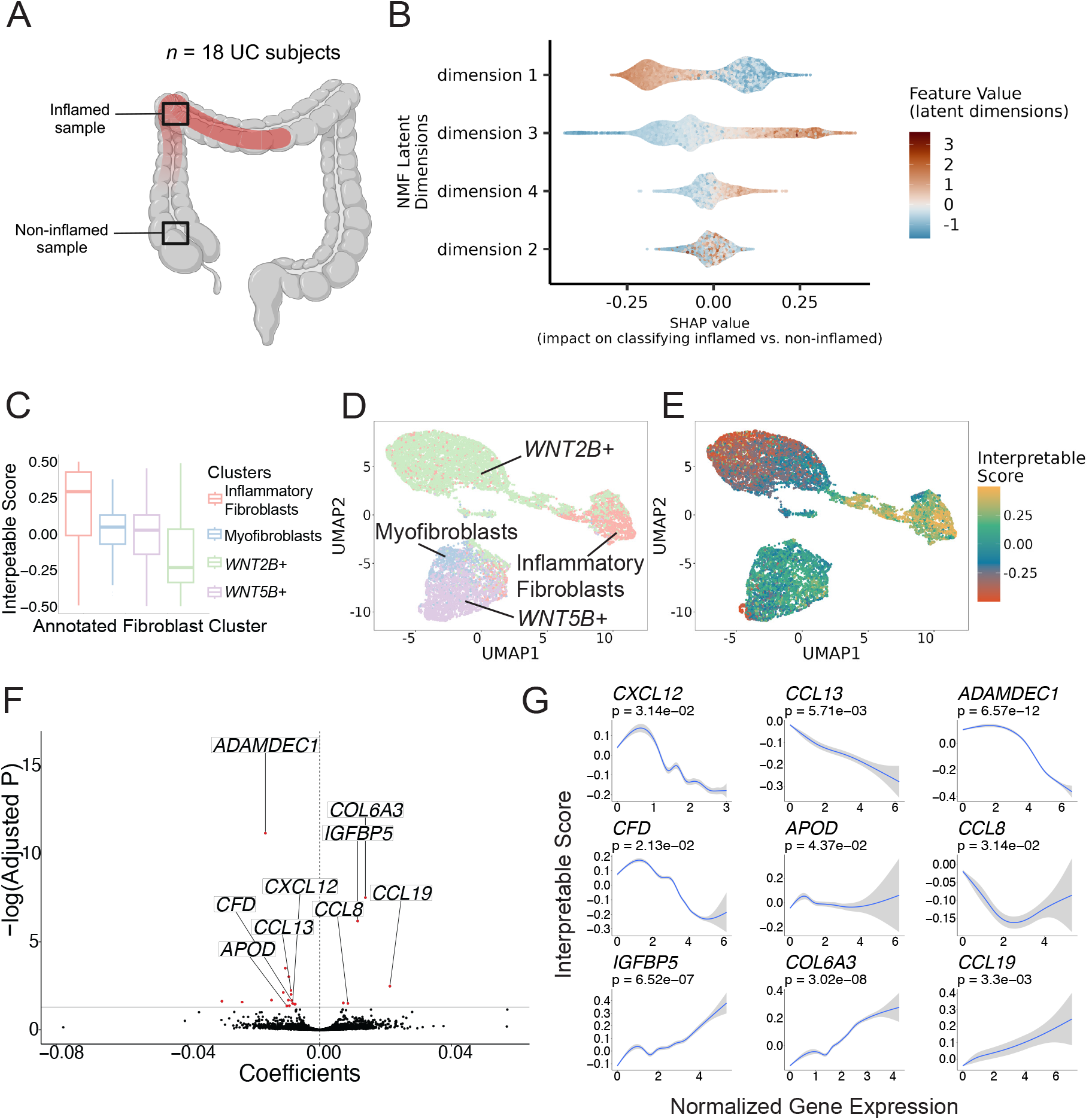
Applying CellPhenoX to single-cell fibroblast transcriptomics from ulcerative colitis tissue. **A**. Schematic depicting the study design, **B**. SHAP summary plot showing the NMF latent dimensions ordered by the mean absolute SHAP value, where each point represents an individual cell, the color denotes the feature value, and the position along the x-axis represents the corresponding SHAP value, **C**.Boxplots showing the distribution of the CellPhenoX Interpretable Score across fibroblast clusters annotated by the original study, **D**. UMAP plot colored the annotated fibroblast clusters, **E**. UMAP plot colored by the CellPhenoX Interpretable Score, **F**. Volcano plot displaying the correlation coefficients between each gene’s expression and the Interpretable Score on the x-axis, with the adjusted p-value on the y-axis. Red points indicate genes with significant correlations, **G**. Correlation plots of normalized gene expression (x-axis) and Interpretable Score (y-axis) for selected significant genes, including 95% confidence intervals and corresponding p-values.

## DISCUSSION

Here we introduced CellPhenoX, a computational method designed to identify cell phenotype shifts by providing interpretable scores that robustly distinguish clinical phenotypes for single-cell omics, including transcriptomics, proteomics, and beyond. Leveraging machine learning techniques and statistical modeling, CellPhenoX uncovers biologically meaningful axes, such as fibroblast differentiation gradients in chronic inflammatory tissues. Intriguingly, a standout function of CellPhenoX is its unique ability to identify cell populations influenced by interaction effects, such as sex and age—an area often overlooked by existing single-cell methods. For example, CellPhenoX successfully linked disease severity with sex and age interactions in simulation and COVID-19 datasets, respectively. This unique capability positions CellPhenoX as an essential tool for uncovering subtle yet significant interaction effects, shedding light on the intricate mechanisms underlying disease pathogenesis.

Moreover, CellPhenoX’s framework integrates flexible decomposition methods including orthogonal and more general matrix decomposition techniques, with adaptable classification algorithms, allowing it to address complex individual-level attributes. One novelty of CellPhenoX is its integration of interpretable AI, which enhances the ability to decipher and link specific cell phenotypes to clinical outcomes with improved explainability and prediction. By leveraging this methodological versatility and interpretability, we demonstrate CellPhenoX’s capability to adapt to diverse clinical contexts, showing its ability to capture nuanced cellular behaviors that could inform clinical phenotyping and potential therapeutic targeting in future.

We noted several limitations and opportunities for CellPhenoX. First, in scenarios where single-cell data display high heterogeneity due to clinical variables overlapping with factors like tissue type or cohort difference, the observed cell abundance shifts associated with certain clinical phenotypes may introduce confounding. While CellPhenoX accommodates random and fixed covariates, it may experience reduced power or potential overfitting in such complex scenarios. Second, we may reveal multiple low-dimensional cell embeddings as interpretable features contributing to the model simultaneously. To maintain the nuance of original characterization, we explain and evaluate the significance of each embedding regarding their contribution to the clinical classifier, and provide a way to aggregate signals to offer global interpretation. Third, applying CellPhenoX on large-scale single-cell datasets may be computational demanding due to the exponential complexity of the Shapley value calculations at individual cell level. As interpretable machine learning advances, we expect to enhance the scalability of CellPhenoX by adapting it to more computationally efficient implementations in the future. Finally, we utilized tree-based models like random forests as they are well-suited for Shapley explanations, maintaining unbiased baseline and marginal Shapley values, while deep learning models usually exhibit biases due to the lack of model-specific algorithms for estimating conditional Shapley values11. As the theory for explainable deep learning advances, we anticipate exploring additional classification methods; however, this is beyond the scope of this work.

In summary, we believe that CellPhenoX is a reliable framework that integrates interpretable machine learning and statistical approaches, an essential milestone as linking single-cell biological insights to clinical impact becomes increasingly critical. While we applied CellPhenoX to single-cell transcriptomics and proteomics, its design—centered on cell lineages, abundance shifts, and interaction effects—should generalize well to other single-cell modalites, such as single-cell ATAC-seq and spatial transcriptomics. Given the unmet need for translating single-cell molecular and cellular findings into clinical applications, such as predicting drug response, tracking disease progression, and identifying sex-based interaction effects, we anticipate CellPhenoX will be valuable for ongoing single-cell translational research in immune-mediated diseases, inflammatory disorders, cancers, and beyond.

## RESOURCE AVAILABILITY

### Lead contact

Information and requests should be directed to the lead contact, Fan Zhang.

### Materials availability

This study did not generate new unique reagents.

### Data and code availability

1. The original data used in this paper can be accessed at the following repositories with details described in the Methods section: 1. Ulcerative Colitis single-cell tissue dataset (Single Cell Portal: SCP259), 2. COVID-19 single-cell PBMC dataset (https://www.ebi.ac.uk/biostudies/arrayexpress/studies/E-MTAB-10026).
2. The CellPhenoX software and source code are available at https://github.com/fanzhanglab/pyCellPhenoX. The single-cell data simulation code is available at https://github.com/fanzhanglab/SCORPIO.
3. Any additional information required to reanalyze the data reported in this paper is available from the lead contact upon request.

## ACKNOWLEDGEMENTS

We thank the members in the Zhang lab for helpful discussions and feedback.

## AUTHOR CONTRIBUTIONS

F.Z and J.Y. conceptualized the study. J.Y. developed and implemented the algorithm, and generated the results and figures. J.I. implemented the single-cell simulation pipeline. R.K. and Z.T.C. finalized the software package. J.Y. and F.Z wrote the manuscript with input from the remaining authors.

## DECLARATION OF INTERESTS

The authors declare no competing interests.

## SUPPLEMENTAL INFORMATION

Document S1. Figures S1-3.

## FIGURE LEGENDS

**Supplementary Figure 1. The simulation schema produces differential abundance signals that influence the model performance**. **A**. Density plots reveal the changes in distribution of the PC loadings by disease status for simulation dataset 1 (top) and simulation dataset 2 (bottom), **B**. Random Forest model performance for simulation dataset 1 (top) and 2 (bottom), including the AUROC curves for different CV repeats (left), the AUPRC curves (middle), and histograms showing the predicted probabilities for the positive (disease) and negative (control) classes.

**Supplementary Figure 2. Details of benchmarking with CNA and MiloR using single-cell simulation datasets**. **A**. UMAP colored by CNA local correlation values (FDR <0.05) for disease for simulation dataset 1, **B**. UMAP colored by CNA local correlation values (FDR <0.05) for simulation dataset 2. Note that cells no passing FDR<0.05 are shown in white, **C**. UMAP colored by MiloR log fold change for simulation dataset 1, **D**. UMAP colored by MiloR log fold change for simulation dataset 2, **E**. UMAP colored by MiloR log fold change by varying the sample proportion parameter value, prop=0.1 (left), prop=0.3 (middle), prop=0.5 (right) (simulation dataset 1), **F**. Boxplots depicting the proportion of correct predictions by CellPhenoX for each simulated cluster. Different thresholds for binarizing the Interpretable Score are applied to facilitate comparison with the simulated disease labels, the top 75th, 80th and 85th percentile of scores are coded as “disease", **G**. Boxplots depicting the proportion of correct predictions by MiloR. Based on the design of the MiloR log fold change metric, thresholds of 2, 4, and 6 are used to evaluate the predicted disease labels for comparison with simulated labels, **H**. Barplots showing the proportion of correct predictions by CNA. In CNA, cells passing an FDR of 0.05 are considered to be significantly associated, which is used herein to compare with the simulated disease status labels

**Supplementary Figure 3. SHAP summary of the analysis for simulated data with interaction effects**. **A**. SHAP summary plot showing the principal components (PCs) and PC:sex interaction terms ranked by mean absolute SHAP value. Each point represents an individual cell, with color indicating the original feature value and position along the x-axis representing the corresponding SHAP value.

**Supplementary Figure 4. Further analytical details of the single-cell ulcerative colitis dataset**. **A**. SHAP summary plot showing the principal components (PCs) ranked by mean absolute SHAP value, where each point represents an individual cell, the color denotes the feature value, and the position along the x-axis represents the corresponding SHAP value (left); UMAP plot colored by the CellPhenoX Interpretable Score (right). **B**. Random Forest performance for the NMF model described in **Figure 4**, including the AUROC curves for the different CV repeats (left), the AUPRC curves (middle), and histograms showing the predicted probabilities for the positive (inflamed) and negative (non-inflamed) classes.

## METHODS

### Datasets Simulation datasets

#### Simulating strategy for single-cell data with differential cluster abundance between conditions

To evaluate the performance of CellPhenoX in identifying association with clinical variables, we developed a custom, flexible simulation pipeline. This approach generates single-cell datasets along with corresponding metadata—including disease status, cell type, subject, batch, as well as age and sex when necessary—specifically tailored to evaluate CellPhenoX. This approach is able to create simulated single-cell cell data, whose feature values represent variance attributable to each factor including cell type clusters and other sample level categorical values based on a Gaussian Mixture Model. This versatile approach allows us to control the source of variation in the dataset through parameters that dictate the proportion of variation explained by each category.

For the analyses in this study, we chose to emphasize the differential abundance of disease-associated cells with respect to cell type clusters. Here, we simulated 30 subjects, each with 100 cells per cell type cluster. Specifically, the simulated dataset included 10 clusters (A through J), where the first seven represented major or more abundant cell types, and the last three were relatively rare cell types. Clusters A and J were designed to be statistically significant expansion in disease compared to control. To achieve differential abundance, we tuned the fold change parameter, which controls the magnitude of increase in the number of disease cells relative to control cells. High fold change values result in more distinct differences in cell proportions between case and control. For the simulation dataset with desired interaction effect, we further manipulated the cell abundance by introducing a conditional abundance influenced by a covariate (e.g., sex) to simulate a condition-biased disease status. This simulation framework provides a robust method for evaluating clinically relevant differential abundance and ensures flexibility in generating datasets reflective of varying biological scenarios, such as rare disease-associated pathogenic cell types.

### Real datasets

#### Ulcerative colitis single-cell tissue dataset

The Ulcerative colitis (UC) single-cell transcriptomics dataset includes cells from healthy (n=12) participants and UC patients (n=18). Two colon biopsies were collected from UC patients, one from inflamed tissue and one from adjacent non-inflamed tissue. We started our analysis using their QC-ed cell matrix, where cluster labels are available driven by their canonical cell lineage marker annotations. Here, we renalyzed the post-QC fibroblasts obtained from the original study, and focused on comparing the inflamed (18 samples, 5,965 cells) and non-inflamed (18 samples, 8,857 cells) gut to identify fibroblast-specific lineages that distinguish these two statuses. The post-QC count matrix, cell type annotation, and sample-level metadata were downloaded from link or database ID (Single Cell Portal: SCP259)24.

#### COVID-19 single-cell PBMC dataset

The COVID-19 dataset is a single-cell peripheral blood mononuclear cells (PBMCs) multi-omics, including surface protein and transcriptome data. Participants in this study ranged the spectrum of disease severity from healthy (24 individuals), hospitalized non-COVID-19 (5 individuals), and intravenous lipopolysaccharide (IV-LPS) (12 individuals) controls, to asymptomatic (12 individuals), mild (26 individuals), moderate (32 individuals), severe (15 individuals), and critical (17 individuals) COVID-19. We obtained the single-cell count matrix together with classified cell type labels, where there were 781,123 total cells after quality control with manually annotated cell populations based on expression of canonical markers and surface proteins. Clinical data is also available, including days from onset, smoker status, age, sex, status on day of sample collection, and the most severe status that the participant progressed to. For our study of interest, we hone in on the single-cell proteomic data from mild (13,590 cells, 26 samples), moderate (19,846 cells, 32 samples), and healthy control (9,692 cells, 24 samples). We downloaded this dataset from the following link: (https://www.ebi.ac.uk/biostudies/arrayexpress/studies/E-MTAB-10026)16.

### Data preprocessing and dimensionality reduction

For single-cell datasets, we first normalized the raw count matrices based on log transformation, followed by selecting highly variable genes based on dispersion. Let this normalized matrix, *Z*, have *M* cells and *P* features. For the *N* samples, we denote *C*(*n*) as the set of cells that come from sample *n*. Anchoring at each cell, we can define a neighborhood based on the adjacency suggested by a nearest neighbor graph. In single-cell transcriptomics data, the neighborhood captures transcriptional similarities, while in single-cell proteomics data, it reflects proteomic similarities. Therefore, two cells that are in close proximity in the neighborhood can be linked by a random walk in the graph. Then, we can generate *A*, a weighted *M* × *M* adjacency matrix that measures the similarity between cells *m* and *m*’ in the graph. In *A*, a number of steps, *s*, and a binary encoded vector, *e*, where the *m*-th entry is 1 and 0 otherwise, we can define the probability that a random walk from *m*’ ends at *m* as

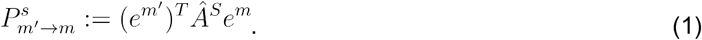

Then, we generate a neighborhood abundance matrix (NAM)9 by first defining

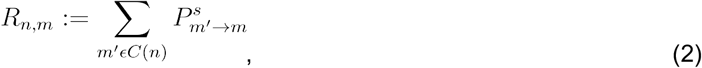

where *Rn,m* is the total number of cells from the *n*-across samples and clinical th sample that are expected to be in neighborhood *m*. Now the NAM, *Q*, is determined by normalizing the rows of *R* so that each row sums to 1 as

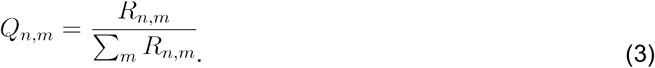

We aim to decrease the number of dimensions of the NAM, *Q*, and then use the derived latent dimensions that summarize the major variation as features to prevent overfitting of the classification model in the next step. Herein, CellPhenoX offers the flexibility to employ different dimensionality reduction techniques on the calculated NAM. Thus, we provide users with the option to choose between two widely used methods for single-cell analyses—principal component analysis (PCA) and non-negative matrix factorization (NMF). A notable difference between these two techniques is that PCA learns orthogonal latent dimensions based on the captured variation, whereas NMF, based on topic modeling, does not enforce orthogonality in the learned dimensions. Specifically, for PCA analyses, we started with learning 100 components, and then selected the top 20 components that achieved the desired variance explained. For NMF, if the number of ranks (cell clusters) to learn, *k*, is not provided by prior knowledge or the users, we tested a range of *k* values and selected the one with the highest silhouette score. In the case of the ulcerative colitis single-cell dataset, we used this as an example to evaluate the performance of both methods, and found that the most influential latent dimensions from both methods summarize similar sources of variation. Considering the event of strong batch effects in single-cell datasets, we applied Harmony19 with default parameters to remove sample-specific and technical batch-associated effects.

### Classification model training and prediction

After dimensionality reduction, we denote *Xi* as the learned latent dimension embeddings, *γ* be the vector(s) of covariates with length *M* (number of cells), *δi* be the variables that measure the interaction effect with latent embedding *i*, and *Y* be the vector for the target variable (clinical phenotype of interest). Herein, *Y* and *γ* are label encoded if they are categorical variables. We can represent the set of all contributing factors, or predictors, as

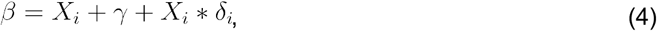

then, we design our classification model, with *θ* parameters, to predict 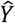 in the following formula

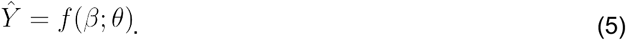

For a random forest model with a categorical outcome, we can further represent 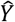 as a majority vote from the *T* trees:

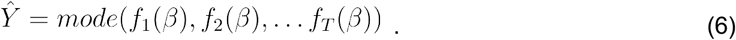

To train the random forest model, we split the data into training, testing, and validation sets using stratified *k*-fold strategy to maintain the relative class proportions in each fold. The outer loop consists of training and testing, while the inner loop involves training and validation. Predictions are made on the testing data from the outer loop, and final model performance is evaluated on the validation set.

### Hyperparameter optimization for classification

To ensure the robustness of our classification model and the resulting SHAP values, we use a nested cross-validation procedure that includes both hyperparameter tuning and cross-validation. Selecting appropriate hyperparameters for the dataset and performing cross-validation to mitigate fold-split bias are crucial for optimal model performance. Since there are many parameters to tune for random forest, we opted for a randomized search across the parameter space instead of an exhaustive search.

### SHAP value calculation

The SHAP value for each feature *βi* is denoted as *Φi*, representing the contribution of *Xi* and other involved covariates to the predicted outcome *Y*. This contribution is determined by

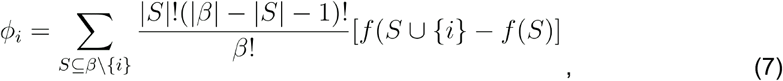

where *S* is a subset of features excluding *βi*, and *f*(*S ∪* {*i*}) and *f*(*S*) are the model predictions with and without *βi*, respectively. In equation (7), we iterate over all possible subsets of features not containing *βi* and calculate the average marginal contribution of each feature *βi* across these subsets. The numerator determines the product of the number of ways to order features in subset *S* and the number of ways to order the remaining features, divided by the total number of ways to order all features in *β*. The average is then multiplied by the difference in model prediction with and without *βi*, highlighting the impact of *βi* on the model output.

For implementation, we adapt the Python shap package’s TreeExplainer function using the best estimator and the testing set for the outer loop. After completing all CV repeats, we iterate over each of the cell IDs and take the average of all SHAP values for each feature across repeats. The resulting data frame maintains the original input dimensions: *N* cells by *M* features. Let *D*% be the data frame containing the SHAP values, where *b* is the number of CV repeats. For each cell *n* we calculate

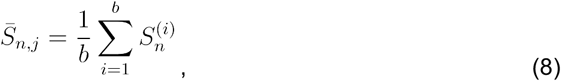

where *S*(*i*) is the *i*-th SHAP matrix,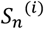 represents the values in row *n*, and *Sn,j* is the mean value at row *n* across all *b* matrices. We then create the final matrix of SHAP values as

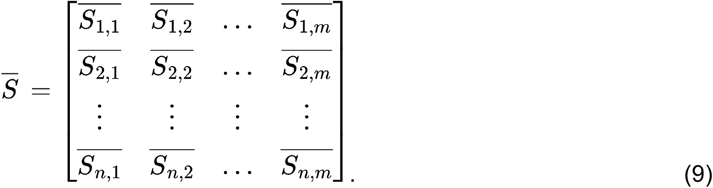

### Multiclass SHAP value aggregation

When dealing with more than two classes, we need to combine the SHAP values from each class to yield one SHAP value per cell per feature. Where there are several ways to do this, this is how we approach it. Let *D* be the matrix of SHAP values with *N* cells and *M* features. For each cell *i*, we select the SHAP values from the matrix that correspond to the cell’s predicted class:

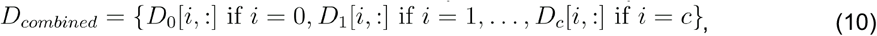

where *c* is the number of classes minus one.

### CellPhenoX Interpretable Score

For each cell *m*, we obtain the Interpretable Score, *ψm*, by summing the obtained SHAP values across feature *i* in *Φm,i*. This score summarizes the contribution of each cell to the overall classification outcome:

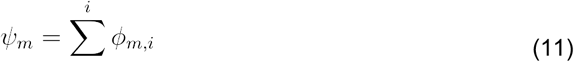

### Benchmarking methods and performance evaluation

*CNA*: CNA (co-varying neighborhood analysis)9 is built to identify co-varying cell abundance shifts across involved individuals with respect to clinical outcomes or sample differences in a linear-based framework. Anchoring at the cell, CNA constructs a nearest neighbor graph describing the transcriptional relatedness of individual cells from each sample. Neighborhoods are determined by quantifying the probability that a random walk from one cell will end at another cell. Then, CNA constructs a neighborhood abundance matrix (NAM), *Q*, a sample by neighborhood matrix, to indicate the relative abundance of neighborhood *m* in sample *n*. Assuming a Gaussian distribution, CNA tests the association with a sample-level covariate or outcome with a linear regression model

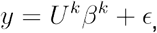

where *UK* denotes the first *K* columns of *Q*’s left singular vectors, *U, βk* is the *K*-length vector of coefficients, and *ϵ* is the zero mean Gaussian noise. A multivariate F-test is employed to determine the P-value across a range of *K* values. Then, given a false discovery rate threshold, the differentially abundant neighborhoods are identified by computing a smoothed correlation between each neighborhood m and the sample-level covariate *y*. We ran CNA with the default parameters given by the tutorials for the R implementation (version 0.0.99), and controlled for sample-sample differences. Specifically, we controlled the sample as covariate from the simulation disease-control single-cell data, and reported cells whose neighborhood correlation to disease at FDR < 0.05.

*MiloR*: MiloR is a method designed to refine the Cydar4 framework by incorporating neighborhood-based analysis for differential abundance8. Milo was chosen for its ability to detect cell population shifts in single-cell data, addressing the limitations of hypersphere-based methods such as Cydar. Milo uses a dynamic k-nearest neighbors (kNN) graph, enabling adaptive and context-sensitive neighborhood formation. The neighborhood formed for a cell is based on its nearest neighbors using the findKNN() function from the BiocNeighbors package. To detect differential abundance, MiloR applies negative binomial generalized linear models (NB-GLMs) via the edgeR package26. To optimize computational efficiency, we first set the parameter prop to 0.1, suggested as default by the developers, to identify cell neighborhoods based on a randomly subsampling. We further performed sensitivity analysis by changing this parameter across 0.1, 0.3, and 0.5. For reproducibility and consistency with established workflows, we implemented MiloR using version 2.1.0 of its R package (https://github.com/MarioniLab/miloR).

## Funding

This work is supported by the PhRMA Foundation, Arthritis National Research Foundation, and the Translational Research Scholars Program award (F.Z.). We also acknowledge support from the NIH NLM Grant T15LM009451 (J.Y.); the Uehara Memorial Foundation Postdoctoral Fellowship, Grant-in-Aid for JSPS Overseas Research Fellows, and the Mochida Memorial Foundation for Medical and Pharmaceutical Research (to J.I.); the Interdisciplinary Biology Program at the Biofrontiers Institute, University of Colorado Boulder, and the National Science Foundation NRT Integrated Data Science Fellowship (award 2022138) (to Z.C.).

## Notes

### Competing Interest Statement

The authors have declared no competing interest.

