## Supplementary figures and images for "CellPhenoX: An eXplainable Cell-specific machine learning method to predict clinical Phenotypes using single-cell multi-omics"

### Supplementary Figures

Supplementary Figure 1

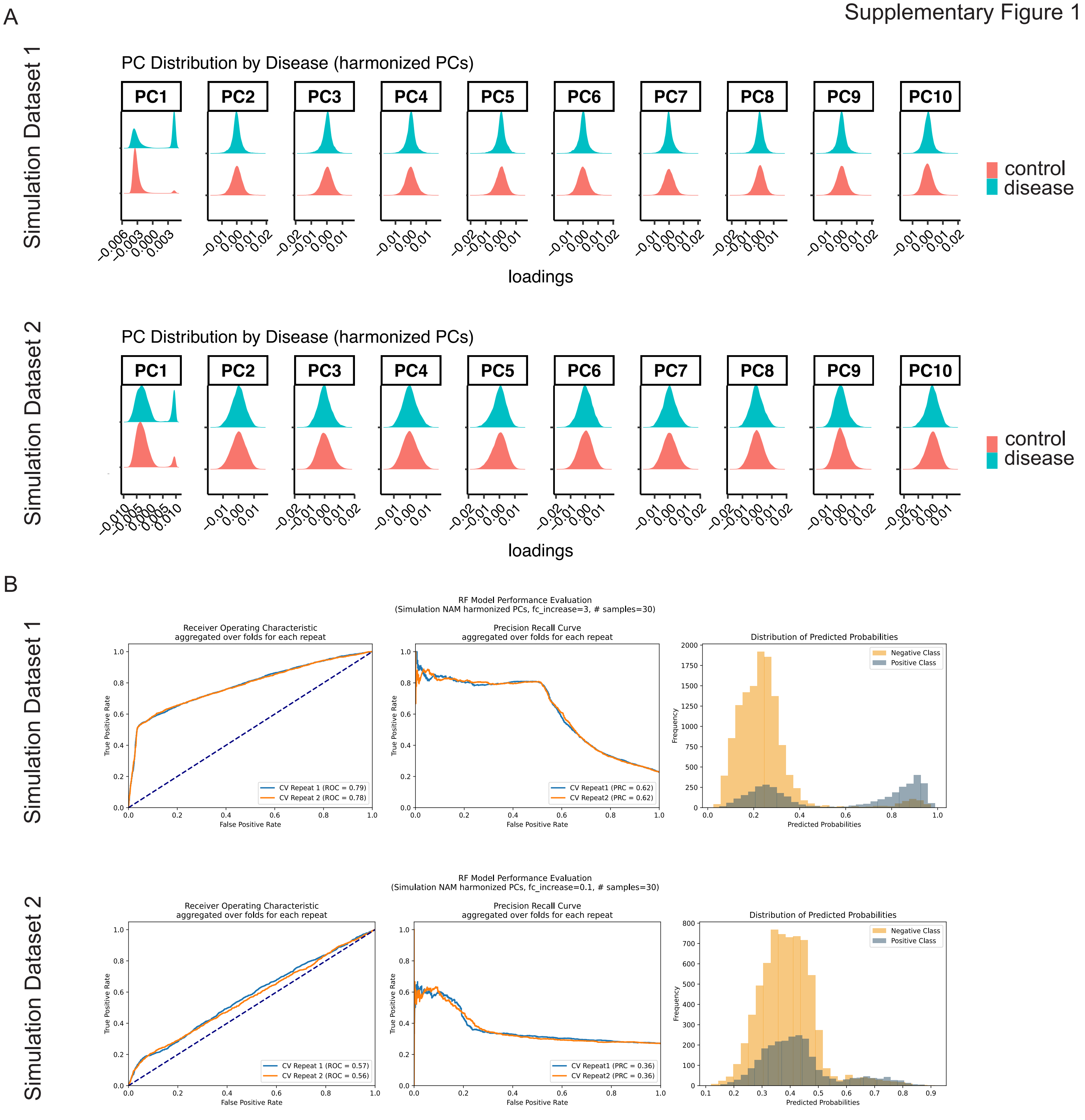

Supplementary FIgure 2

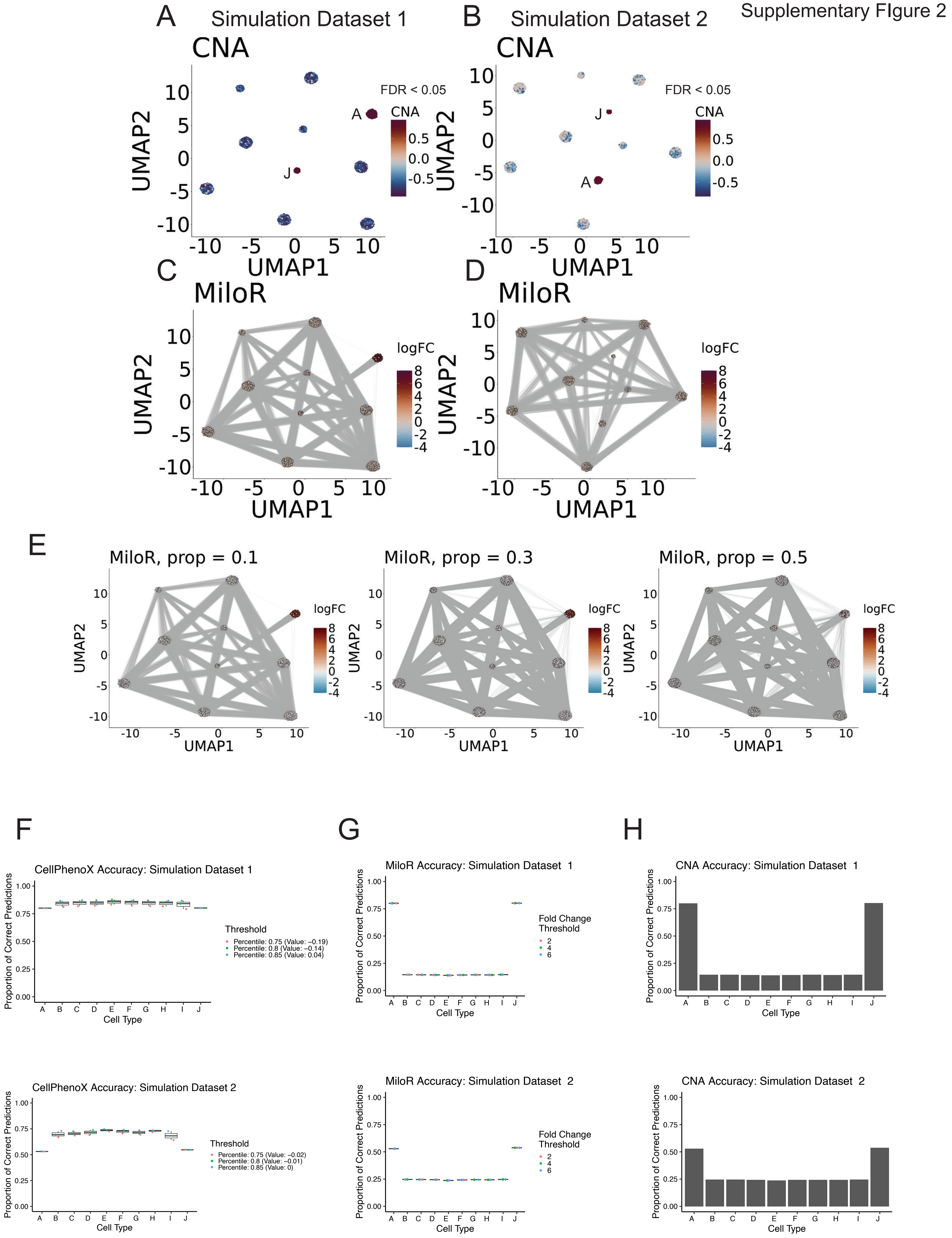

Supplementary Figure 3

A

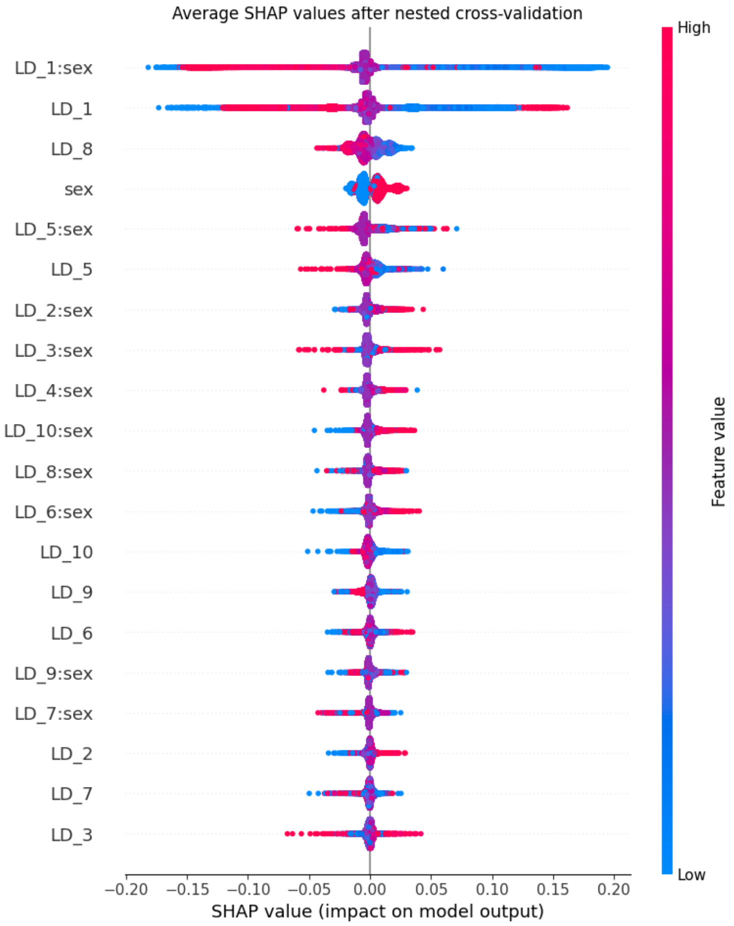

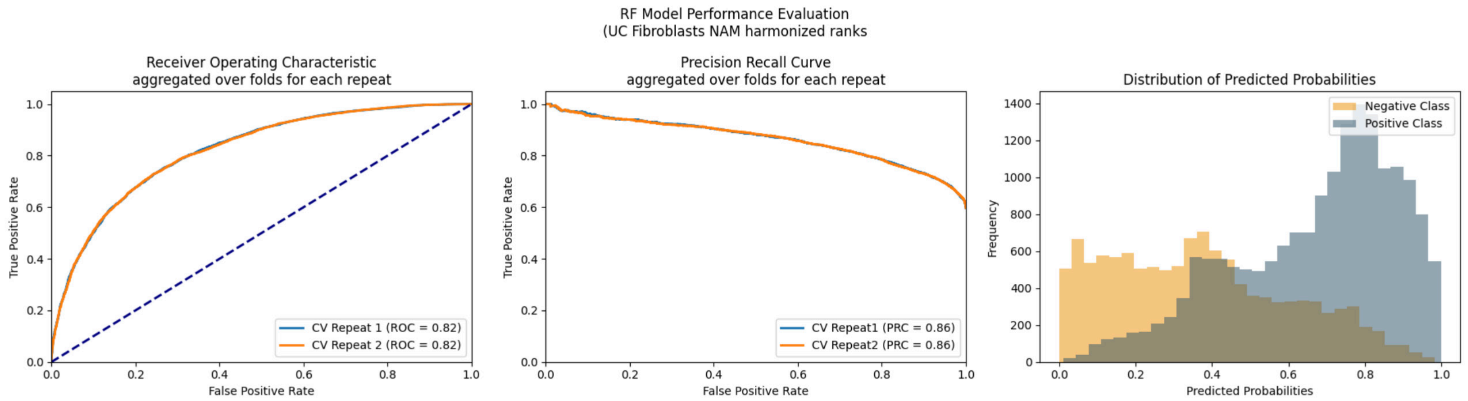

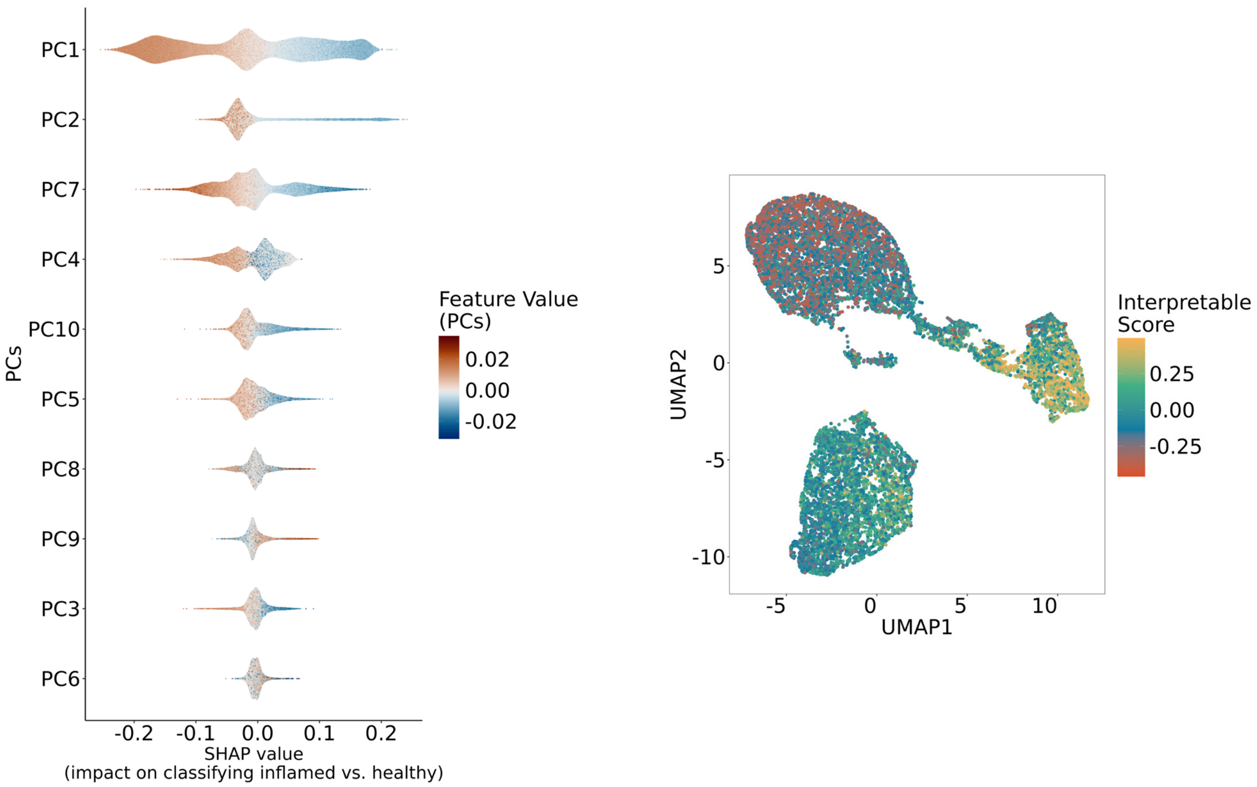
